# The scent of the fly

**DOI:** 10.1101/206375

**Authors:** Paul G. Becher, Sebastien Lebreton, Erika A. Wallin, Erik Hedenström, Felipe Borrero, Marie Bengtsson, Volker Jörger, Peter Witzgall

**Author notes:** Correspondence: Peter Witzgall, SLU, Box 102, 23053 Alnarp, Sweden. Paul G. Becher -, Sebastien Lebreton -, Erika A. Wallin -, Erik Hedenström -, Felipe Borrero -, Marie Bengtsson -, Volker Jörger.

## Abstract

(*Z*)-4-undecenal (*Z*4-11Al) is the volatile pheromone produced by females of the vinegar fly *Drosophila melanogaster*. Female flies emit *Z*4-11Al for species-specific communication and mate-finding. A sensory panel finds that synthetic *Z*4-11Al has a characteristic flavour, which we perceive even at the small amounts produced by one female fly. Since only females produce *Z*4-11Al, and not males, we can reliably distinguish between single *D. melanogaster* males and females, according to their scent. Females release *Z*4-11Al at 2.4 ng/h and we readily sense 1 ng synthetic *Z*4-11Al in a glass of wine (0.03 nmol/L), while a tenfold concentration is perceived as a loud off-flavour. This corroborates the observation that a glass of wine is spoilt by a single *D. melanogaster* fly falling into it, which we here show is caused by *Z*4-11Al. The biological role of *Z*4-11Al or structurally related aldehydes in humans and the basis for this semiochemical convergence remains yet unclear.

## Introduction

All living things communicate with chemicals. Unlike sounds or sights, semiochemicals interconnect species across the kingdoms, and enable information exchange between animals, plants and microorganisms (Schultz and Appel 2004). A fascinating, recurrent observation is that the same compound is bioactive in different species and context. Evolutionary convergence may result from the widespread occurrence or even physico-chemical properties facilitating information transmission, but is first of all thought to reflect the biological significance of chemicals, including the underlying biochemical pathways and precursors.

Citrus fruit is a preferred oviposition substrate for the vinegar fly *Drosophila melanogaster* (Dweck et al. 2013) provided that yeast is present (Becher et al. 2012). Citrus peel and brewer’s yeast both produce linalool (Chisholm et al. 2003; Carrau et al. 2005), which the flies perceive via several odorant receptors (Ors), including DmelOr69a (Münch et al. 2016; Lebreton et al. 2017).

Linalool, commonly found in the headspace of foliage, flowers and fruit, is bioactive in many animals. Plant-produced linalool enhances mate-finding in several phytophagous insects, while other species release linalool as a sex pheromone component (Hefetz et al. 1979; Aldrich et al. 1986; Leal et al. 1993; Yang et al. 2004). Herbivory, on the other hand, upregulates linalool production in plants, which may protect against further infestation (Mithöfer and Boland 2012). The (R) and (S) enantiomers of linalool differentially attract pollinators and herbivores, for feeding and oviposition (Reisenman et al. 2010; Saveer et al. 2012; Raguso 2016), and enantiomeric changes during phenological development modulate our perception of flower aroma (Pragadheesh et al. 2017). In mammals, linalool induces psychopharmalogical effects via glutamate receptors (Elisabetsky et al. 1995; Nakamura et al. 2009), perception via Ors produces a sweet, floral note and makes a prominent contribution to the bouquet of flowers, fruit and wine, where both grape and yeast are a source of linalool (Lewinsohn et al. 2001; Carrau et al. 2005; Swiegers et al. 2005).

The response to food and mate olfactory cues is strongly interconnected in *Drosophila* (Lebreton et al. 2015; Gorter et al. 2016; Das et al. 2017) and DmelOr69a is also tuned to the newly identified female pheromone (*Z*)-4-undecenal (*Z*4-11Al), in addition to food odorants (Lebreton et al. 2017). Curiously, Z4-11Al is also found in citrus essential oil (Chisholm et al. 2003). Unsaturated aldehydes are prominent constitutents of a range of food aromas, including fruit, wine, meat and fish (e.g. Varlet et al. 2006; Cullere et al. 2007, Perez-Cacho and Rouseff 2008; Shi et al. 2013; Yang et al. 2008). Moreover, Z4-11Al is an anal gland volatile in the rabbit (Goodrich et al. 1978).

While collecting volatiles from *D. melanogaster* flies, we discovered that we can reliably distinguish single male from female flies by their scent, which is strongly reminiscent of synthetic *Z*4-11Al. We then employed a sensory panel to verify whether we can indeed discern single flies, and whether the newly discovered pheromone *Z*4-11Al contributes to the scent of the female fly.

## Materials and methods

### Chemicals

Isomeric and chemical purity of synthetic *Z*4-11Al were 98.6% and >99.9%, respectively, according to gas chromatography coupled to mass spectrometry (6890 GC and 5975 MS, Agilent Technologies, Santa Clara, CA, USA). Ethanol (redistilled, >99.9% purity; Merck, Darmstadt, Germany) was used as solvent.

### Sensory evaluation

Eight members (six men and two women) of the sensory panel for organoleptic tests for the wine-growing area of Baden evaluated the odor of *D*. *melanogaster* and synthetic *Z*4-11Al. Members of this panel have been trained and selected for the official quality assessment of wines produced in Baden, at the Federal Institute for Viticulture, Freiburg, Germany. Each test comprised three glasses, control and two treatments, which were presented in random order. The panel was asked to score odor intensity, ranging from 1 (weak, silent) to 9 (strong, loud) and to comment on odor quality. The local human subjects committee approved sensory evaluation of Z4-11Al by sniffing. The first test compared the odor from single male and female flies. Flies were kept during 5 min in empty wine tasting glasses (215 ml) and were released shortly before tests. The second test compared a glass impregnated with fly odor and *Z*4-11Al (10 ng in 10 µl ethanol), which was applied to an empty glass, the solvent was allowed to evaporate during 2 min. Next, 10 ng *Z*4-11Al or a female fly were added to a glass filled with either water or white wine (dry Pinot blanc, Freiburg 2013, Staatsweinkellerei Freiburg). The fly was removed after 5 min, prior to testing. Finally, 1 or 5 ng *Z*4-11Al was added to wine.

### Statistical analysis

Odor panel data was analyzed using one-tailed analysis of variance (ANOVA) followed by a Tukey test. Normality was tested using Shapiro-Wilk and homoscedasticity was tested using Levene’s test. All analysis were carried out using SPSS v. 20 (IBM Corp, 2011).

## Results

*D. melanogaster* females (Fig. 1) produce a distinctive scent. The sensory panel found the odor of single female flies to be stronger and qualitatively clearly different from male flies (Fig. 2a).

**Fig.1.**
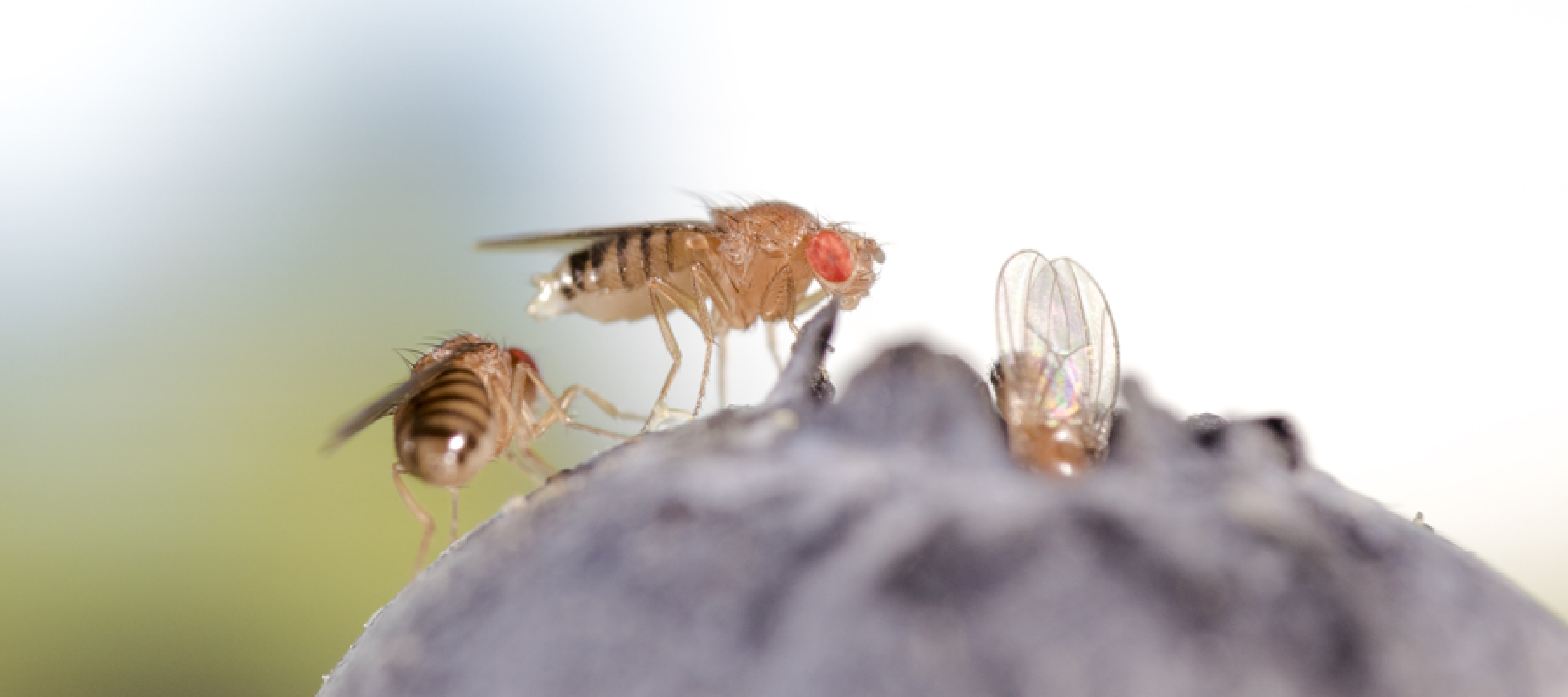
Fruit fly *D. melanogaster* female with exposed ovipositor on blueberry (Photo by Cyrus Mahmoudi).

**Fig.2.**
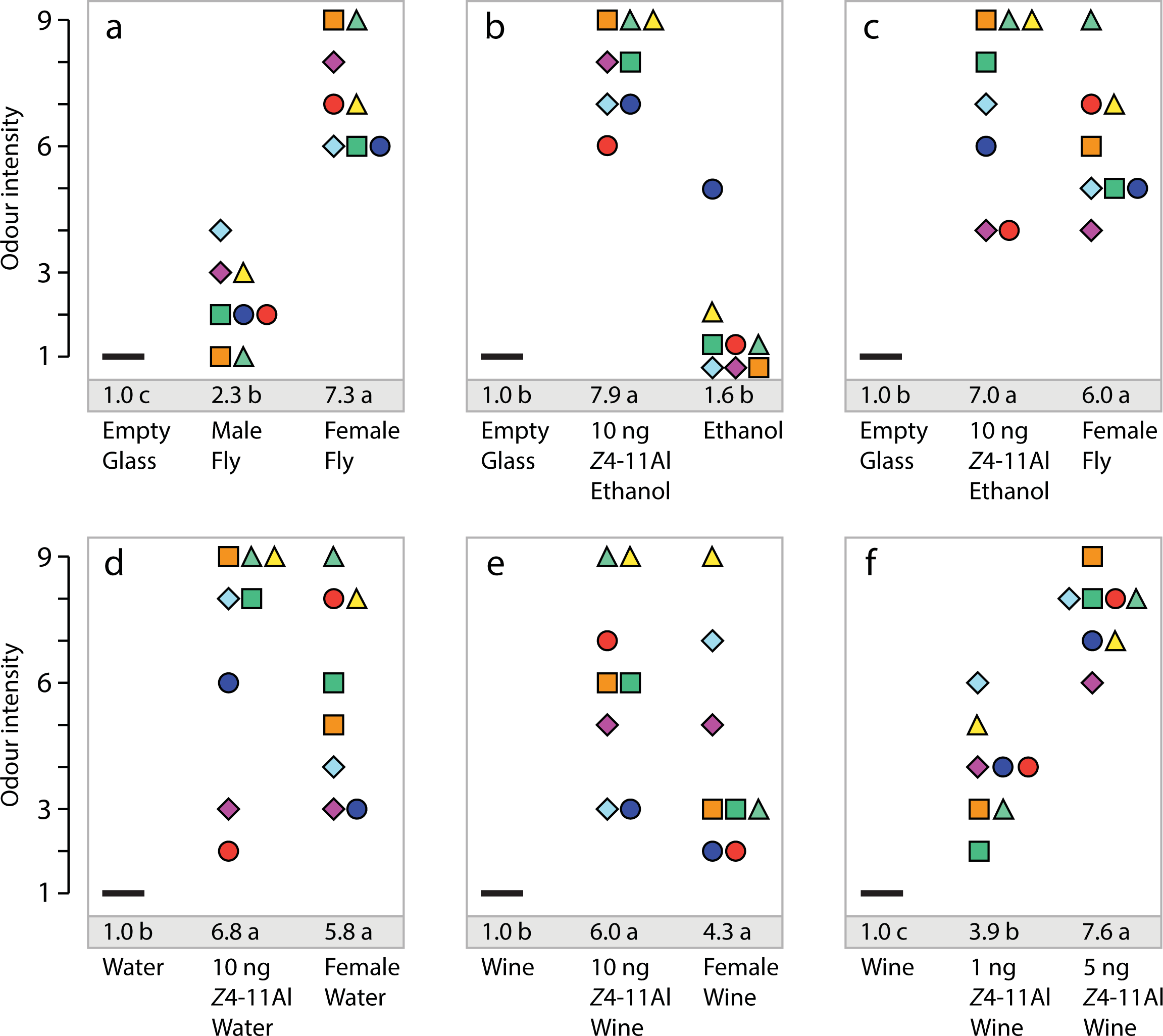
Sensory evaluation of fly odor and synthetic (*Z*)-4-undecenal (*Z*4-11Al). Odor intensity scale ranges from 1 (weak) to 9 (strong), symbols show evaluation by individual test panel members, mean intensity ratings followed by different letters are significantly different (p<0.001). Olfactory intensity of (a) the odor of a single *D. melanogaster* male and female fly adsorbed during 5 min in an empty wine glass (F=96.711), (b) 10 ng synthetic Z4-11Al and solvent (ethanol) (F=106.732), (c) 10 ng Z4-11Al and the odor of a single *D. melanogaster* female fly in an empty glass (F=34.720), (d) in a glass with water (F=16.689), (e) in a glass with wine (F=12.952), (f) 1 ng and 5 ng Z4-11Al in a glass with wine (F=110.694).

Chemical analysis has shown earlier that *Z*4-11Al and its precursor, the cuticular hydrocarbon (*Z,Z*)-7,11-heptacosadiene, are produced by female flies, not by males (Billeter et al. 2009; Lebreton et al. 2017). Our panel tests established that synthetic *Z*4-11Al has a distinctive odor (Fig. 2b). Moreover, a female fly and 10 ng *Z*4-11Al were found to be similar, with respect to odor quality and intensity, when presented in an empty glass, in water or wine (Fig. 2c,d,e). Since 10 ng *Z*4-11Al was assessed as slightly louder than the odor of a fly, we compared *Z*4-11Al at 1 ng and 5 ng, showing that as little as 1 ng *Z*4-11Al in a wine glass (corresponding to ca. 5 ng/L or 0.03 nmol/L at the time of application) was clearly perceptible (Fig. 2f). Even at small amounts, *Z*4-11Al was perceived as a somewhat unpleasant off-flavour.

The detection threshold for *Z*4-11Al is apparently similar in flies and men, since we clearly sense *Z*4-11Al released from a single fly (Fig. 2a). Chemical analysis found that *D. melanogaster* females released *Z*4-11Al at a rate of 2.4 ng/h and solvent extracts of fly cuticula contained 0.3 ng *Z*4-11Al/female (Lebreton et al. 2017).

## Discussion

Sensory evaluation confirmed that we sensitively smell *Z*4-11Al, the female-produced pheromone of the fruit fly *D. melanogaster* (Lebreton et al. 2017) and that we reliably distinguish single female from male flies. This supports the observation that one fly spoils a glass of wine, after falling into it - provided it is of the female sex. Other fly volatiles may contribute to our perception of fly odor. However, *Z*4-11Al is the most abundant compound which is released by females only, other volatiles are found in both sexes (Lebreton et al. 2017).

A straightforward explanation for convergent perception of *Z*4-11Al is, however, not at hand. The occurrence of *Z*4-11Al in nature is probably incompletely known and its possible role in humans remains unclear. *Z*4-11Al may merely be reminiscent of other food aldehydes, or it might avert ingestion of fruit infested with vinegar flies, which may be contaminated with microbes vectored by flies.

A range of mammalian Ors is responsive to aldehydes (e.g. Benbernou et al. 2007; Saito et al. 2009; Nara et al. 2011), including human OR1A1 and OR2W1 (Geithe et al. 2017a,b). Perception of *Z*4-11Al effects the heart rate of the rabbit, where it has been found in the anal gland (Goodrich et al. 1978). The positional isomer (*E*)-2-undecenal is a bovid body odor (Gikonyo et al. 2002) and olfactory sensory neurons of ticks (Acari, Ixodidae) tuned to aldehydes afford indirect evidence for aldehyes as vertebrate signals (Steullet and Guerin 1994). A characteristic scent wich is reminiscent of tangerine emanates from colonies of crested auklet, a monogamous seabird. Two unsaturated aldehydes, the chain-shortened analog (*Z*)-4-decenal (*Z*4-10Al) and (*Z*)-2-decenal are main odor-active constitutents. In crested auklet, *Z*4-10Al likely plays a role as an ectoparasite repellent and a signal of mate quality (Douglas et al. 2001; Hagelin et al. 2003; Caro and Balthazart 2010).

*Z*4-11Al is also part of clementine aroma (Chisholm et al. 2003) and it may play a dual role as social signal and food cue, not only in flies, but also in other animals. The olfactory sense in animals plays a key role during habitat adaptation. Tuning of Ors to habitat cues will create a bias for mate-finding signals that match or are structurally similar to habitat odorants (Endler 1992). This idea yields a tentative scenario for the convergence of semiochemicals. Insects and other animals feed on fruit, containing associated yeasts that facilitate digestion of plant materials, provide nutrients and protection of food from antagonistic microorganisms. Animal-produced compounds sharing structural motifs may have secondarily been adopted as mating signals, via established sensory channels dedicated to habitat and food odorants.

The olfactory system of *Drosophila* is conveniently accessible to experimental investigation and current research extends beyond the Or ligand repertoire (Münch and Galizia 2016) to neural circuits underlying odor-mediated behavior (Kohl et al. 2013; Auer and Benton 2016; Seki et al. 2017), chemical ecology (Depetris-Chauvin et al. 2015; Mansourian and Stensmyr 2015) and phylogenetic diversification (Shiao et al. 2015; Arguello et al. 2016; Ramasamy et al. 2016). A future challenge is to extend functional, behavioral, ecological and phylogenetic studies to include vertebrates, towards an understanding of the chemical vocabulary that interconnects us with other living things.

## Acknowledgements

We thank the members of the sensory panel (Freiburg, Germany) for evaluating the fly scent and Cyrus Mahmoudi (comgraphix.de, Germany) for sharing a fruit fly photograph. Supported by the Linnaeus initiative “Insect Chemical Ecology, Ethology and Evolution” (Formas, SLU).

## Data accessibility

Data is completely included in the figure.

## Authors’ contributions

P.G.B. sensed the fly scent, P.G.B., S.L., M.B. and V.J. conceived the idea and contributed to the experiment, F.B. calculated statistics, E.W. and E.H. synthesized the test chemical, P.W. supervised the project and wrote the manuscript, all authors contributed to and approved the final version of the manuscript.

## References

Aldrich JR, Lusby WR, Kochansky JP (1986) Identification of a new predaceous stink bug pheromone and its attractiveness to the eastern yellowjacket. Cell Molec Life Sc 42:583–585. (doi:10.1007/BF01946714)

Arguello JR, Cardoso-Moreira M, Grenier JK, Gottipati S, Clark AG, Benton R (2016) Extensive local adaptation within the chemosensory system following *Drosophila melanogaster*’s global expansion. Nature Comm 7:11855 (doi:10.1038/ncomms11855)

Auer TO, Benton R (2016) Sexual circuitry in *Drosophila*. Curr Op Neurobiol 38:18–26. (doi:10.1016/j.conb.2016.01.004)

Becher PG, Flick G, Rozpedowska E, Schmidt A, Hagman A, Lebreton S, Larsson MC, Hansson BS, Piskur J, Witzgall P, Bengtsson M (2012) Yeast, not fruit volatiles mediate attraction and development of the fruit fly *Drosophila melanogaster*. Funct Ecol 26:822–828 (doi:10.1111/j.1365-2435.2012.02006.x)

Benbernou N, Tacher S, Robin S, Rakotomanga M, Senger F, Galibert F (2007) Functional analysis of a subset of canine olfactory receptor genes. J Heredity 98:500–505. (doi:10.1093/jhered/esm054)

Billeter JC, Atallah J, Krupp JJ, Millar JG, Levine JD (2009) Specialized cells tag sexual and species identity in *Drosophila melanogaster*. Nature 461:987–U250. (doi:10.1038/nature08495)

Caro SP, Balthazart J (2010) Pheromones in birds: myth or reality? J Comp Physiol A 196:751–766. (doi:10.1007/s00359-010-0534-4)

Carrau FM, Medina K, Boido E, Farina L, Gaggero C, Dellacassa E, Versini G, Henschke PA (2005) De novo synthesis of monoterpenes by *Saccharomyces cerevisiae* wine yeasts. FEMS Microbiol Lett 243:107–15. (doi:10.1016/j.femsle.2004.11.050)

Chisholm MG, Jell JA, Cass DM. (2003) Characterization of the major odorants found in the peel oil of *Citrus reticulata* Blanco cv. Clementine using gas chromatography-olfactometry. Flavour Fragrance J 18:275–281. (doi:10.1002/ffj.1172)

Cullere L, Cacho J, Ferreira V (2007) An assessment of the role played by some oxidation-related aldehydes in wine aroma. J Agric Food Chem 55:876–881. (doi:10.1021/jf062432)

Das S, Trona F, Khallaf MA, Schuh E, Knaden M, Hansson BS, Sachse S. 2017. Electrical synapses mediate synergism between pheromone and food odors in *Drosophila melanogaster*. Proc Natl Acad Sc USA 114:9962–9971. (doi:10.1073/pnas.1712706114)

Depetris-Chauvin A, Galagovsky D, Grosjean Y (2015) Chemicals and chemoreceptors: ecologically relevant signals driving behavior in *Drosophila*. Front Ecol Evol 3:41. (doi:10.3389/fevo.2015.00041)

Douglas H, Jones T, Conner W (2001) Heteropteran chemical repellents identified in the citrus odor of a seabird (crested auklet: *Aethia cristatella*): evolutionary convergence in chemical ecology. Naturwissensch 88:330–332. (doi:10.1007/s001140100236)

Dweck HK, Ebrahim SA, Kromann S, Bown D, Hillbur Y, Sachse S, Hansson BH, Stensmyr MC (2013) Olfactory preference for egg laying on citrus substrates in *Drosophila*. Curr Biol 23:2472–80. (doi: 10.1016/j.cub.2013.10.047)

Elisabetsky E, Marschner J, Souza DO (1995) Effects of linalool on glutamatergic system in the rat cerebral cortex. Neurochem Res 20:461–465. (doi:10.1007/BF00973103)

Endler JA (1992) Signals, signal conditions, and the direction of evolution. Am Naturalist 139:S125–S153. (doi:10.1086/285308)

Geithe C, Noe F, Kreissl J, Krautwurst D (2017a) The broadly tuned odorant receptor OR1A1 is highly selective for 3-methyl-2, 4-nonanedione, a key food odorant in aged wines, tea, and other foods. Chemical Senses 42:181–193. (doi:10.1093/chemse/bjw117)

Geithe C, Protze J, Kreuchwig F, Krause G, Krautwurst D (2017b) Structural determinants of a conserved enantiomer-selective carvone binding pocket in the human odorant receptor OR1A1. Cell Molec Life Sci 74:4209–4229. (doi:10.1007/s00018-017-2576-z)

Gikonyo NK, Hassanali A, Njagi PG, Gitu PM, Midiwo JO (2002) Odor composition of preferred (buffalo and ox) and nonpreferred (waterbuck) hosts of some savanna tsetse flies. J Chem Ecol 28:969–981. (doi:10.1023/A:1015205716921)

Goodrich BS, Hesterman ER, Murray KE, Mykytowycz R, Stanley G, Sugowdz G. (1978) Identification of behaviorally significant volatile compounds in the anal gland of the rabbit, *Oryctolagus cuniculus*. J Chem Ecol 4:581–594. (doi:10.1007/BF00988922)

Gorter JA, Jagadeesh S, Gahr C, Boonekamp JJ, Levine JD, Billeter JC (2016) The nutritional and hedonic value of food modulate sexual receptivity in *Drosophila melanogaster* females. Sci Rep 6:19441. (doi:10.1038/srep19441)

Hagelin JC, Jones IL, Rasmussen LEL (2003) A tangerine-scented social odour in a monogamous seabird. Proc R Soc B 270:1323–1329. (doi: 10.1098/rspb.2003.2379)

Hefetz A, Batra SWT, Blum MS (1979) Linalool, neral and geranial in the mandibular glands of *Colletes* bees - an aggregation pheromone. Cell Molec Life Sc 35:319–320. (doi:10.1007/BF01964324)

Kohl J, Ostrovsky AD, Frechter S, Jefferis GS (2013) A bidirectional circuit switch reroutes pheromone signals in male and female brains. Cell 155:1610–1623. (doi:10.1016/j.cell.2013.11.025)

Leal WS, Sawada M, Matsuyama S, Kuwahara Y, Hasegawa M (1993) Unusual periodicity of sex pheromone production in the large black chafer *Holotrichia parallela*. J Chem Ecol 19:1381–1391. (doi:10.1007/BF00984883)

Lebreton S, Trona S, Borrero-Echeverry F, Bilz F, Grabe V, Becher PG, Carlsson MA, Nässel DR, Hansson BS, Sachse S, Witzgall P (2015) Feeding regulates sex pheromone attraction and courtship in *Drosophila* females. Sci Rep 5:13132 (doi:10.1038/srep13132)

Lebreton S, Borrero-Echeverry F, Gonzalez F, Solum M, Wallin E, Hedenström E, Hansson BS, Gustavsson A-L, Bengtsson M, Birgersson G, Walker WB, Dweck H, Becher PG, Witzgall P (2017) A *Drosophila* female pheromone elicits species-specific long-range attraction via an olfactory channel with dual specificity for sex and food. BMC Biology 15:88. (doi:10.1186/s12915-017-0427-x)

Lewinsohn E, Schalechet F, Wilkinson J, Matsui K, Tadmor Y, Nam KH, Amar O, Lastochkin E, Larkov O, Ravid U, Hiatt W, Gepstein S, Pichersky E (2001) Enhanced levels of the aroma and flavor compound S-linalool by metabolic engineering of the terpenoid pathway in tomato fruits. Plant Physiol 127:1256–1265. (doi:10.1104/pp.010293)

Mansourian S, Stensmyr MC (2015) The chemical ecology of the fly. Curr Op Neurobiol 34:95–102. (doi:10.1016/j.conb.2015.02.006)

Mithöfer A, Boland W (2012) Plant defense against herbivores: chemical aspects. Annu Rev Plant Biol 63:431–450. (doi:10.1146/annurev-arplant-042110-103854)

Münch D, Galizia CG. (2016) DoOR 2.0 - comprehensive mapping of *Drosophila melanogaster* odorant responses. Sci Rep 6:21841. (doi:10.1038/srep21841)

Nakamura A, Fujiwara S, Matsumoto I, Abe K (2009) Stress repression in restrained rats by (R)-(–)-linalool inhalation and gene expression profiling of their whole blood cells. J Agric Food Chem 57:5480–5485. (doi:10.1021/jf900420g)

Nara K, Saraiva LR, Ye X, Buck LB (2011) A large-scale analysis of odor coding in the olfactory epithelium. J Neurosc 31:9179–9191. (doi:10.1523/jneurosci.1282-11.201)

Perez-Cacho PR, Rouseff RL (2008) Fresh squeezed orange juice odor: a review. Critical Rev Food Sci Nutr 48:681–695. (doi: 10.1080/10408390701638902)

Pragadheesh VS, Chanotiya CS, Rastogi S, Shasany AK (2017) Scent from *Jasminum grandiflorum* flowers: investigation of the change in linalool enantiomers at various developmental stages using chemical and molecular methods. Phytochemistry 140:83–94. (doi:10.1016/j.phytochem.2017.04.018)

Raguso RA (2016) More lessons from linalool: insights gained from a ubiquitous floral volatile. Curr Op Plant Biol 32:31–36. (doi:10.1016/j.pbi.2016.05.007)

Ramasamy S, Ometto L, Crava CM, Revadi S, Kaur R, Horner DS, Pisani D, Dekker T, Anfora G, Rota-Stabelli O (2016) The evolution of olfactory gene families in *Drosophila* and the genomic basis of chemical-ecological adaptation in *Drosophila suzukii*. Gen Biol Evol 8:2297–2311. (doi:10.1093/gbe/evw160)

Reisenman CE, Riffell JA, Bernays EA, Hildebrand JG (2010) Antagonistic effects of floral scent in an insect-plant interaction. Proc R Soc B 277:2371–2379. (doi:10.1098/rspb.2010.0163)

Saito H, Chi Q, Zhuang H, Matsunami H, Mainland JD (2009) Odor coding by a mammalian receptor repertoire. Science Signal 2(60):ra9. (doi:10.1126/scisignal.2000016)

Saveer AM, Kromann S, Birgerson G, Bengtsson M, Lindblom T, Balkenius A, Hansson BS, Witzgall P, Becher PG, Ignell R (2012) Floral to green: mating switches moth olfactory coding and preference. Proc R Soc B 279:2314–2322. (doi:10.1098/rspb.2011.2710)

Schultz JC, Appel HM (2004) Cross-kingdom cross-talk: hormones shared by plants and their insect herbivores. Ecology 85:70–77. (doi:10.1890/02-0704)

Seki Y, Dweck HK, Rybak J, Wicher D, Sachse S, Hansson BS (2017) Olfactory coding from the periphery to higher brain centers in the *Drosophila* brain. BMC Biol 15:56 (doi:10.1186/s12915-017-0389-z)

Shi X, Zhang X, Song S, Tan C, Jia C, Xia S (2013) Identification of characteristic flavour precursors from enzymatic hydrolysis-mild thermal oxidation tallow by descriptive sensory analysis and gas chromatography-olfactometry and partial least squares regression. J Chromatogr B 913:69–76. (doi:10.1016/j.jchromb.2012.11.032)

Shiao MS, Chang JM, Fan WL, Lu MY, Notredame C, Fang S, Kondo R, Li WH (2015) Expression divergence of chemosensory genes between *Drosophila sechellia* and its sibling species and its implications for host shift. Gen Biol Evolution 7:2843–2858. (doi:10.1093/gbe/evv183)

Steullet P, Guerin PM (1994) Identification of vertebrate volatiles stimulating olfactory receptors on tarsus I of the tick *Amblyomma variegatum* Fabricius (Ixodidae). 1. Receptors within the Haller’s organ capsule. J Comp Physiol A 174:27–38. (doi:10.1007/BF00192003)

Swiegers JH, Bartowsky EJ, Henschke PA, Pretorius IS (2005) Yeast and bacterial modulation of wine aroma and flavour. Austral J Grape Wine Res 11:139–173. (doi:10.1111/j.1755-0238.2005.tb00285.x)

Varlet V, Knockaert C, Prost C, Serot T (2006) Comparison of odor-active volatile compounds of fresh and smoked salmon. J Agric Food Chem 54:3391–3401. (doi:10.1021/jf053001p)

Yang Z, Bengtsson M, Witzgall P (2004) Host plant volatiles synergize response to sex pheromone in codling moth, *Cydia pomonella*. J Chem Ecol 30:619–629. (doi:10.1023/B:JOEC.0000018633.94002.af)

Yang DS, Shewfelt RL, Lee KS, Kays SJ (2008) Comparison of odor-active compounds from six distinctly different rice flavor types. J Agric Food Chem 56:2780–2787. (doi:10.1021/jf072685t)

